# Baicalein ameliorates experimental ulcerative colitis recurrency by downregulating neonatal Fc receptor via the NF-κB signaling pathway

**DOI:** 10.1101/2024.10.22.619759

**Authors:** Haoyang Hu, Fuliang Lu, Xudong Guan, Xuehua Jiang, Chengming Wen, Ling Wang

## Abstract

Ulcerative colitis (UC) is a chronic autoimmune disease (AID) that causes mild to moderate, unpredictable symptoms, including diarrhea and abdominal pain. Against neonatal Fc receptors (FcRn) has been proven a unique AID treatment strategy by decreasing the effects of pathogenic autoantibody. Our previous study revealed that FcRn inhibition is beneficial in UC treatment through reducing colonic neutrophil extracellular traps (NET) formation via accelerating serum anti-neutrophil cytoplasm antibodies (ANCA) clearance. In this study, we initially confirmed the specific impact of downregulating FcRn in preventing UC relapse by injecting rAAV, that carrying Fcgrt-shRNA, in mice. Next, we investigated the inhibition effects and regulation mechanisms of baicalein (BCL) on FcRn and assessed its capacity to withstand UC recurrence using NCM460 cells and dextran sodium sulfate-induced mice models by determining the expression of FcRn and its related transcription factors. We also measured colonic NET-associated protein (NAP) expression and serum concentrations of IgG, ANCA, TNF-α, IL-1β, and c-reactive protein (CRP). UC inflammation severity was determined using the disease activity index (DAI) and histopathological score (HS). BCL treatment remarkably decreased the mRNA and protein contents of FcRn, p50, and p65 but did not impact STAT1 expression and the phosphorylation of IκB and STAT1. Long-term BCL administration inhibited colonic FcRn expression and reduced serum ANCA levels, colonic NAPs expression, serum inflammation-related indexes (including TNF-α, IL-1β, and CRP), DAI, and HS scores in UC mice during inflammation relapse better than salazosulfapyridine. Our study indicates that BCL ameliorates UC recurrency by inhibiting FcRn expression via p50/p65 heterodimer-mediated NF-κB signaling.

**Chemical compounds:** Chemical compounds studied in this article were listed below:

Baicalein (PubChem CID: 5281605); Salazosulfapyridine (PubChem CID: 5339).

## 1. Introduction

Ulcerative colitis (UC), an inflammatory bowel disease[1, 2], is a long-term illness prone to recurrence and canceration[3]. Increasing evidence indicates that UC is induced by an abnormal immune response against luminal antigens that results in persistent and uncontrolled inflammation[4–6]. Neutrophils are the first immune cells recruited to the site of inflammation in areas with mucosal damage from UC[7]. These activated neutrophils promote the resolution of ongoing inflammation and mucosal healing and liberate neutrophil extracellular traps (NET)[8, 9]. NET formation leads to the generation of antineutrophil cytoplasm autoantibodies (ANCA) that can be used as a biomarker to determine UC severity and guide treatment[10, 11]. The in vivo transport of ANCA is mediated by the neonatal Fc receptor (FcRn), that functions within intracellular endosomes where it binds IgG and albumin at distinct and nonoverlapping sites under acidic pH conditions[12, 13]. FcRn-mediated recycling causes circulating IgG and albumin to have a long half-life[14–16]; however, the long half-life of specific autoantibodies and their associated immune complexes can cause pathogenic effects in patients with autoimmune diseases[17, 18].

Baicalein (BCL), a flavonoid derived from *Scutellaria baicalensis* Georgi, displays various pharmacological activities, including anti-inflammatory, antioxidant, and antitumor effects[19–21], which is mainly due to its ability to scavenge reactive oxygen species and improve antioxidant status by attenuating NF-κB activity and suppressing the expression of several inflammatory cytokines and chemokines[22–25]. Several studies have reported that BCL has therapeutic benefits for UC treatment due to its antioxidant and anti-inflammatory activities and its ability to repair the intestinal epithelium[26, 27]; however, the effects of BCL on FcRn and inflammatory recurrence in UC remain unclear.

NF-κB is one of the most studied transcription factors and is crucial in regulating immune response and inflammation[28]. The NF-κB family includes five master transcription factors that regulate the transcription of target genes by binding to specific DNA elements (κB sites) as various hetero-or homo-dimers: NF-κB1 (also named p50); NF-κB2 (also named p52); RelA (also named p65); RelB; and c-Rel[29]. A wide range of stimuli trigger the assembly of multiprotein complexes, thus activates the IκB kinases and consequently mediating NF-κB activation[30]. Previously, Xindong Liu and his colleagues reported that NF-κB signaling via intronic sequences regulates FcRn expression and function; they also indicated that FcRn expression is decreased by IFN-γ via the JAK/STAT1 pathway[31, 32].

Several recent studies have indicated that an FcRn blockade may serve as a unique treatment for IgG-mediated autoimmune diseases[33–37]. Our previous study revealed that the targeted inhibition of FcRn is beneficial in UC treatment due to its ability to reduce colonic NET formation via the acceleration of serum ANCA clearance, especially during the inflammation recurrence period (IRP)[38]. In this study, we aim to explore the effects of BCL on colonic FcRn expression and its regulation mechanism in UC and evaluate the potential usage of BCL in UC recurrency treatment based on FcRn inhibition using the dextran sodium sulfate (DSS)-induced mice model.

## 2. Materials and methods

### 2.1. Materials

The materials used in this study included: DSS (purity > 98 %; Yeasen Biological Technology, Shanghai, China); purified human ANCA protein (purity > 97 %, HENGYUAN Biotechnology, Shanghai, China); Recombinant Human TNF-α Protein (purity > 97 %, ABclonal Technology, Wuhan, China); Recombinant Human IFN-γ Protein (purity > 95 %, ABclonal Technology); BCL (purity > 98 %; Meilun Biological Technology, Dalian, China); Salazosulfapyridine (SASP, purity > 97 %; Meilun Biological Technology); and fecal occult blood test kits (Leagene Biotechnology, Beijing, China).

### 2.2. Cell Culture and experimental design

NCM460 human normal colon epithelial cells were obtained from INCELL (Texas, USA) and cultured at 37°C in a humidified atmosphere containing 5%CO_2_. The cells were routinely screened for mycoplasma contamination. To determine the effects of BCL on FcRn expression in NCM460 cells, we first assessed the cytotoxicity of a series of BCL concentrations (10, 20, 40, 80, or 100 μM) in NCM460 cells after 48 h of culture. The cells were treated with nontoxic concentrations in subsequent experiments. After incubation, total RNA and protein were extracted. The mRNA and protein expression of FcRn were compared to those of the blank control using a real-time polymerase chain reaction (RT-qPCR) and western blotting.

To further evaluate the regulation mechanism of BCL on FcRn, we treated NCM460 cells with BCL at the concentration that had the most significant inhibition effect on FcRn for 48 h with and without small interfering RNA (siRNA)-targeted p50, p65, and STAT1. After incubation, total RNA and protein were extracted. The mRNA and protein expressions of FcRn, p50, p65, IκB, pIκB, STAT1, and phosphorylated STAT1 (pSTAT1) were compared to those of the blank control using RT-qPCR and western blotting. To verify the effects of BCL on TNF-α or IFN-γ activated NF-κB or STAT1 signaling in FcRn transcriptional regulation, we stimulated NCM460 cells with 50 ng/mL TNF-α or IFN-γ, accompanied with BCL treatment at the concentration that had the most significant inhibition effect on FcRn for 48 h with and without small interfering RNA (siRNA)-targeted p50, p65, and STAT1. After incubation, total RNA was extracted, and the mRNA expressions of FcRn were compared to those of the blank control using RT-qPCR.

### 2.3. MTT Assay

MTT assays were performed to evaluate the cytotoxicity of BCL on NCM460 cells. Briefly, NCM460 cells were seeded in a 96-well plate at 4 × 10^4^ cells/well density and cultured in Dulbecco’s Modified Eagle Medium (DMEM) containing 10% fetal bovine serum (FBS). The following day, the culture medium was removed, and 100 µL BCL solution in concentrations of 10, 20, 40, 80, or 100 μM prepared in DMEM containing 1 % FBS or blank DMEM (control) were added to each well. The cells were then incubated at 37°C for 24 h. Next, the BCL solution was removed and 100 µL MTT solution (0.5 mg/mL dissolved in PBS buffer) was added to each well. The plate was then incubated for 4 h at 37°C. After incubation, the medium was removed and 100 µL DMSO was added to the wells to solubilize the formazan product. Finally, a colorimetric assay was performed at 490 nm using a Multiskan MK3 Reader (Thermo Fisher Scientific, Waltham, MA, USA).

### 2.4. In vitro ANCA recycling assay in NCM460 cells

To perform the ANCA recycling assay, NCM460 cells were seeded into 12-well plates at 2 × 10^5^cells/well density and cultured in a DMEM medium containing 10% FBS. When the cells had grown to 95–100% confluency, they were washed and starved for 1 h using Hank’s Balanced Salt Solution (HBSS; pH 7.4). Next, purified human ANCA protein was diluted to 100 ng/mL in HBSS and added to the cells, which were then incubated for 4 h. Afterwards, the medium was removed and the cells were extensively washed with ice-cold HBSS, then incubated with DMEM for 12 h. The medium and cells were then collected to analyze their ANCA concentrations and FcRn expression levels using ELISA and western blotting, respectively.

### 2.5. siRNA knockdown of p50/p65/STAT1 expression in NCM460 cells

NCM460 cells were transfected with a mixture of control siRNAs or p50/p65/STAT1-specific siRNAs (si-p50/si-p65/si-STAT1; Tsingke Biological Technology, Chengdu, China). Each siRNA sequence is provided in Table S4. For each transfection, a siRNA mixture was diluted in the transfection medium, and reagent as described by the manufacturer (Yeasen Biological Technology Co. Ltd., Shanghai, China). The cells were incubated for 24 h at 37°C in a CO_2_ incubator with a siRNA mixture and BCL solution diluted in DMEM without FBS. Next, the incubation solution was replaced with fresh BCL solution diluted in DMEM containing 1 % FBS for another 24 h. After incubation, the total RNA and protein were extracted and the mRNA and the protein expressions of FcRn, p50, p65, and STAT1 were compared with those of the blank control using RT-qPCR and western blotting.

### 2.6. Animals and experimental design

C57BL/6 mice (in-house, random-bred) aged 8–12 weeks and weighing 25–30 g were obtained from Ensiweier Biological Technology (Chongqing, China) and quarantined in the animal house of the West China School of Pharmacy, Sichuan University (Chengdu, China). All animal experiments were conducted according to the guidelines of the National Institutes of Health guide for the care and use of Laboratory animals (NIH Publications No. 8023, revised 1978), and approved by the Animal Ethics Committee of Sichuan University (No. K2022006).

Our previous study demonstrated the efficacy of anti-FcRn therapy in a rat model through the intravenous administration of anti-rat Fc heavy chain heterodimer monoclonal antibodies (anti-Fc-mAb). In this study, we further investigated the impact of reducing colonic FcRn expression on treating UC. For this purpose, we administered Fcgrt-shRNA encapsulated in a recombinant adeno-associated virus (rAAV-Fcgrt-shRNA, provided by Landmbio Co. Ltd, Guangzhou, China) to suppress colonic FcRn expression in mice via intraperitoneal injection. We evaluated the disease activity and colonic histopathological changes in a DSS-induced UC mouse model throughout the experimental inflammation progression and recurrence phase. The specific experimental procedure is illustrated in Supplementary Figure S2.

To evaluate the effects of BCL on FcRn and its regulation mechanism and therapeutic impact on UC recurrence based on FcRn inhibition in UC mice, 54 mice were randomly divided into three groups (n = 18 for each group). The groups were administrated with either ultra-pure (UP) water (UC group), SASP (400 mg/kg, p.o.; SASP group), or BCL (100 mg/kg, p.o.; BCL group) throughout the entire experiment period (day 1–28). In addition, each group received 2.5 % (w/v) DSS for 7 days, then received UP water for 14 days to simulate a remission period. The 2.5 % (w/v) DSS solution was then administered for an additional 7 days to simulate an inflammation recurrence period until the end of the experiment (day 28). Six mice were sacrificed on days 7 and 28. Supplementary data Fig. S3 shows the entire experimental process. Blood samples (500 μL) were collected from the retro-orbital plexus of each mouse and placed into microcentrifuge tubes; then, the mice were killed by cervical dislocation before removing their colons for biochemical and histopathological analyses.

### 2.7. Disease activity index (DAI) and histopathological score (HS) evaluation

The DAI for each mouse was calculated based on body weight loss, occult blood, and stool consistency as described in Table S1 on a scale of 1–3 or 1–4 for each parameter and a total score of 11. The collected colon tissues were fixed in 4% paraformaldehyde, embedded in paraffin, then stained with hematoxylin and eosin. Histological damage was scored based on goblet cell loss, mucosa thickening, inflammatory cell infiltration, submucosa cell infiltration, ulcers, and crypt abscesses, as described in Table S2. A score of 1–4 was given for each parameter, with a maximal total score of 19 for HS.

### 2.8. ELISA

Mice blood samples were centrifuged at 4000 rpm for 10 min to obtain serum. IgG, ANCA, TNF-α, IL-1β, and CRP serum concentrations were measured using the applicable ELISA kits (HENGYUAN Biotechnology, Shanghai, China) according to the manufacturer’s instructions. Mice colons were homogenized in a RIPA lysis buffer (Solarbio, Beijing, China) and centrifuged at 12000 rpm for 10 min to obtain tissue total protein sample. Colonic concentrations of TNF-α and IL-1β were measured using the applicable ELISA kits (HENGYUAN Biotechnology) according to the manufacturer’s instructions.

### 2.9. RT-qPCR analysis

Total RNA extraction from NCM460 cells and mice colons were performed using TRIzol reagent (Gbcbio, Guangzhou, China) according to the manufacturer’s instructions. RNA (1 µg) was used as a template for cDNA synthesis using Hifair™ 1st strand cDNA Synthesis SuperMix (Yeasen Biological Technology Co. Ltd. Shanghai, China). qRT-PCR was performed using Hieff™ qPCR SYBR® Green Master Mix (Yeasen Biological Technology) with specific primers (Table S3). The amplification protocol consisted of initial denaturation at 95°C for 5 min, followed by 40 denaturation cycles at 95°C for 10 s, then annealing at 60°C for 20 s, and extension at 72°C for 20 s. The relative gene expression was normalized against that of human GAPDH or mice gapdh. Gene expression was calculated using the 2^−ΔΔCT^ method. The primers were obtained from Tsingke Biological Technology (Chengdu, China).

### 2.10. Western blot analysis

Cells and mice colons were homogenized in a RIPA lysis buffer (Solarbio). Whole-cell or tissue extracts were prepared by direct lysis in 1 × electrophoresis sample buffer. The protein content was determined using a BCA protein assay kit (Biyuntian Co Ltd., Shanghai, China). Total cellular protein was resolved by 10% SDS-PAGE and transferred onto a polyvinylidene difluoride membrane. The membrane was blocked with 5% nonfat milk and incubated with the primary antibody overnight at 4°C, followed by incubation with the secondary antibody for 1 h. Antibodies against FcRn, p50, p65, IκB, phosphorylated IκB (pIκB), STAT1, phosphorylated STAT1 (pSTAT1), protein arginine deiminase-4 (PAD4), GAPDH and β actin were obtained from ABclonal Technology (Wuhan, China). Antibodies against citrullinated histone H3 (citH3) were obtained from Proteintech Biotechnology (Wuhan, China). All antibodies were used at the dilutions recommended by the manufacturers. Before incubation with the specific primary antibody, membranes were cut horizontally according to each protein’s molecular weight (MW). For proteins with close MW, the same sample would run separately for each protein, such as IκB/pIκB and stat1/pstat1. Blots were observed with ECL luminescence imager (MiniChemi500, Sage Creation Science Co. LTD, Beijing, China). The densities of the protein bands were determined using ImageJ software (version 1.8.0_345, National Institutes of Health, Bethesda, MD, USA).

### 2.11. Statistical analysis

Statistical analysis was performed using IBM SPSS Statistics version 22 (IBM, Armonk, NY, USA). One-way ANOVA with Bonferroni’s multiple comparison test was used to analyze most sets of quantitative data. If the data did not meet normality or homogeneity of variance, nonparametric analysis using the Kruskal–Wallis test was conducted. All other analyses were performed using the Student’s t-test. The level of significance was set at p < 0.05. All data are presented as the mean ± standard deviation.

## 3. Results

### 3.1. Knockdown colonic FcRn expression interrupted ANCA-NET cycle and ameliorated UC relapse in mice

We reported earlier that anti-FcRn therapy has benefits in UC treatment (especially during the IRP) through reducing colonic NET formation by accelerating serum ANCA clearance in rat through intravenous injection of anti-Fc-mAb[38]. In this study, we further investigated the effects of downregulating of colonic FcRn expression with rAAV-Fcgrt-shRNA on UC treatment during the experimental-simulated inflammation progression and recurrence status. The results showed that FcRn mRNA and protein expression was significantly downregulated in group AAV-A or AAV-B compared with group UC-A or UC-B, respectively; However, there was no statistical significance in the changes of FcRn mRNA or protein in group SASP compared with group UC-B. (Fig. 1a-d) Furthermore, it can be seen that the changes in mouse serum IgG levels are basically consistent with the changes in FcRn expression trend (Fig 1e). Additionally, we tracked the serum ANCA levels of mice in each group. As depicted in Figure 1f, the data indicated that injection of rAAV-Fcgrt-shRNA notably decreased serum ANCA levels compared to the UC control group. Furthermore, during IRP, it demonstrated a more potent ANCA elimination capability than SASP administration. Besides, the changes of colonic NET-associated protein (NAP, includes PAD4 and citH3) were in consistent with the ANCA changes in each group (Fig1g and 1h). Collectively, these data elucidated that rAAV-Fcgrt-shRNA injection can directedly reduce colonic FcRn expression, accelerated ANCA clearing from serum and led to the reduction of NET formation in mice colon.

**Fig. 1.**
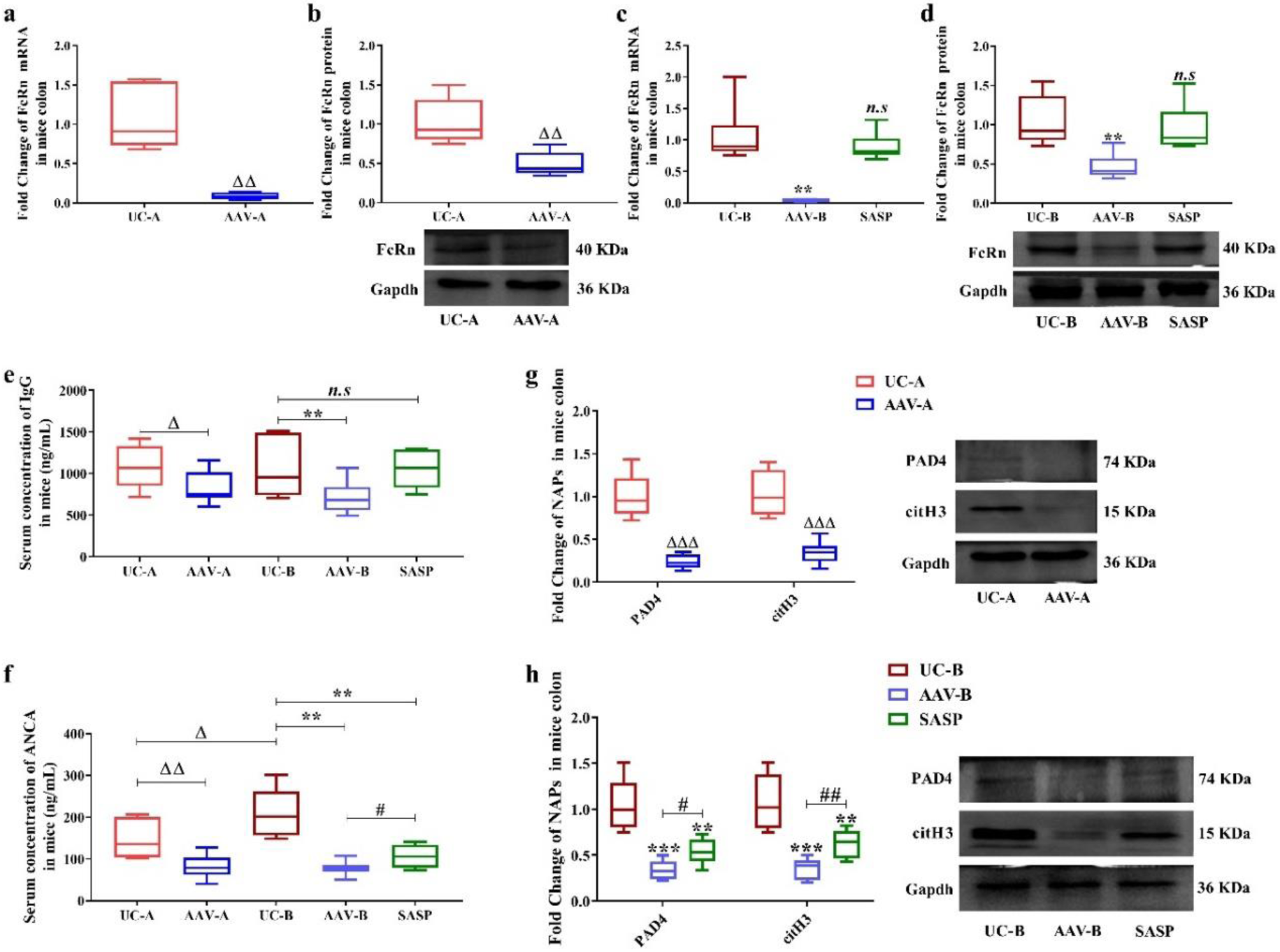
Effects of FcRn knockdown by rAAV-Fcgrt-shRNA on colonic ANCA-NET cycle during inflammation progression and relapse in mice. UC-A and AAV-A indicated mice in each group were sampling from inflammation progression period (noted as N1 shown in Fig.S2); UC-B, AAV-B and SASP indicated mice in each group were sampling from inflammation recurrency period (noted as N2 shown in Fig.S2) (a-d) Colonic FcRn mRNA or protein contents in mice of each group; the protein or mRNA expression levels of FcRn were normalized to those of gapdh. (e, f). serum IgG and ANCA concentrations in mice of each group. (g, h) Colonic PAD4 and citH3 protein contents in mice of each group; the protein expression levels of PAD4 and citH3 were normalized to those of gapdh. All data are expressed as the mean ± S.D. (n = 6 replicates/treatment).Δ/*, ΔΔ/**, and ΔΔΔ/*** represents p < 0.05, p < 0.01 and p < 0.001 against UC-A or UC-B, respectively; #, and ## represents p < 0.05, and p < 0.01 against AAV-B; *n.s* represents the differences that were not statistically significant (p > 0.05) against UC-A or UC-B.

In the follow-up study, we evaluated the UC therapeutic effects of colonic FcRn downregulating during the inflammation progression period and IRP. In particular, the DAI and HS scores (Fig. 2a-b) showed significant reduction in groups AAV-A and AAV-B compared to the UC control. Moreover, levels of the serum inflammation-related index (IRI), such as TNF-α, IL-1β, and CRP (Fig. 2d-f), were markedly decreased in these groups. Among these markers, it was observed that injection of rAAV-Fcgrt-shRNA resulted in lower levels compared to SASP, which aligns with the findings of reduced serum ANCA, and colonic NAPs contents detected earlier. Collectively, these findings once again confirm the efficacy of anti-FcRn therapy in mice with UC. This is achieved by downregulating colonic FcRn expression through the injection of rAAV-Fcgrt-shRNA, resulting in reduced formation of colonic mucosal NETs by enhancing the clearance of serum ANCA.

**Fig. 2.**
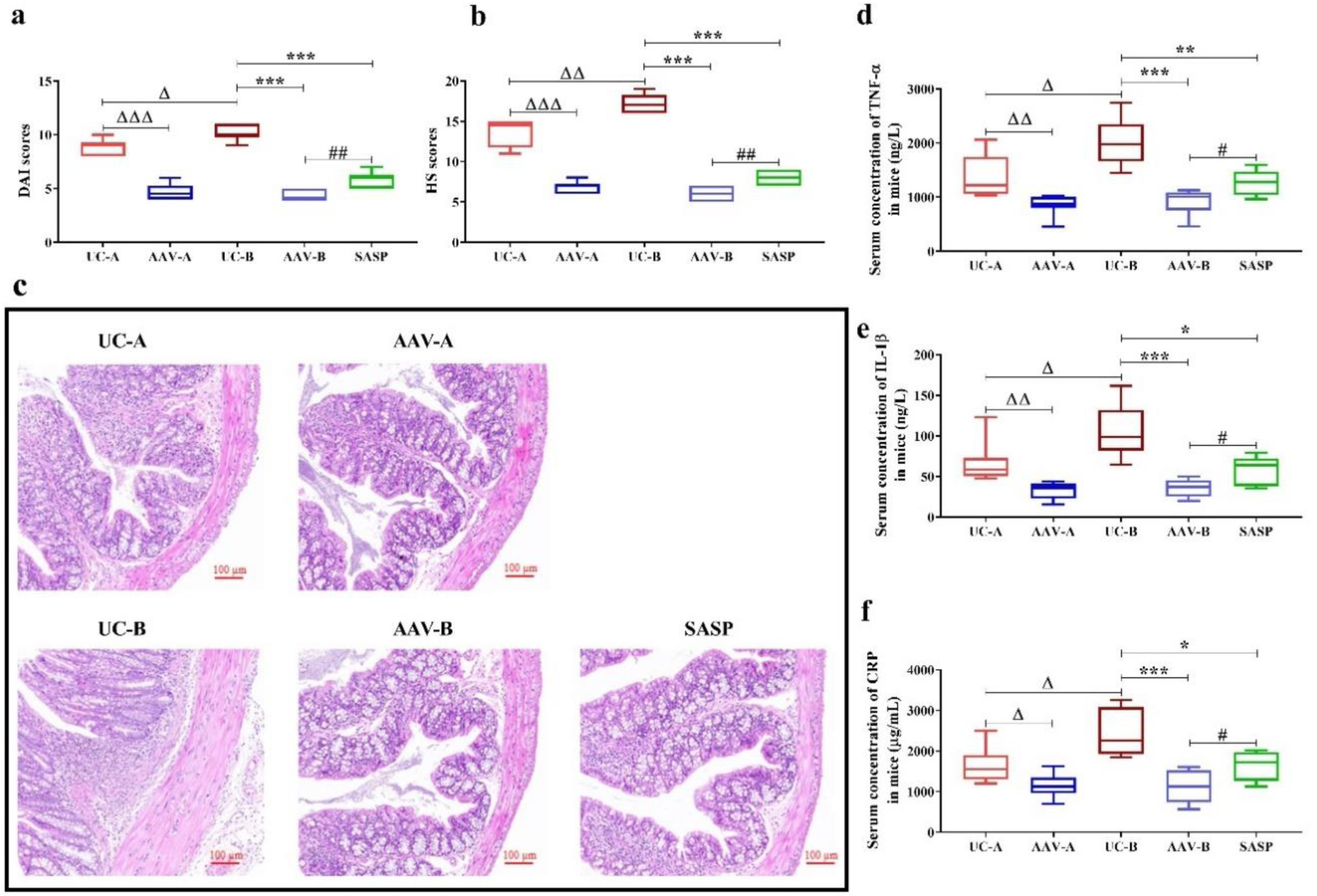
Effects of FcRn knockdown by rAAV-Fcgrt-shRNA on UC inflammation progression and relapse in mice. UC-A and AAV-A indicated mice in each group were sampling from inflammation progression period (noted as N1 shown in Fig.S2); UC-B, AAV-B and SASP indicated mice in each group were sampling from inflammation recurrency period (noted as N2 shown in Fig.S2) (a, b) Calculation of DAI and HS in mice with UC of each group. (c) The representative histopathological images of mice colon in each group. (d–f) Serum concentrations of TNF-α, IL-1β, and CRP in UC mice of each group. All data are expressed as the mean ± S.D. (n = 6 replicates/treatment). Δ/*, ΔΔ/**, and ΔΔΔ/*** represents p < 0.05, p < 0.01 and p < 0.001 against UC-A or UC-B, respectively; #, and ## represents p < 0.05, and p < 0.01 against AAV-B.

### 3.2. BCL inhibits FcRn expression and ANCA recycling in NCM460 cells

We determined that BCL concentrations at 10, 20, and 40 μM were nontoxic to the NCM460 cells (Figure S1) and used these to treat the NCM460 cells to assess the effects of BCL on FcRn expression. We found that 20 or 40 μM BCL significantly reduces the mRNA and protein expression of FcRn compared to the control, with 40 μM BCL exhibiting the most significant inhibition effects. In contrast, 10 μM BCL only inhibited the mRNA expression of FcRn and did not inhibit the protein (Fig. 3 a-b). These results suggest that FcRn inhibition at these concentrations is dose-dependent. Next, we pretreated NCM460 cells with 40 μM BCL and found that ANCA recycling was markedly decreased after BCL treatment (Fig. 3c).

**Fig. 3.**
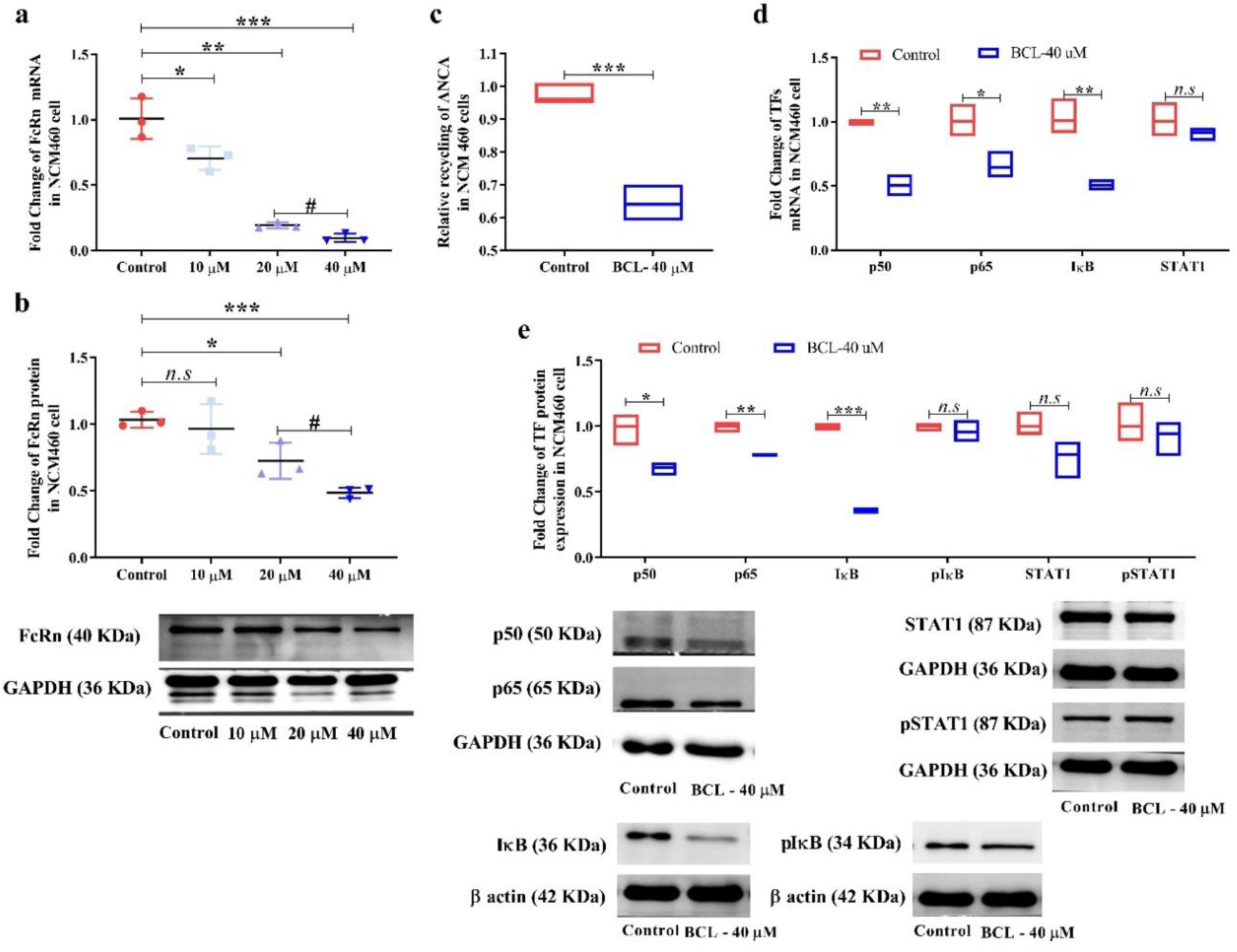
Effects of BCL on FcRn and related transcription factors expression and ANCA recycling in NCM460 cells. (a, b) FcRn mRNA and protein contents in NCM460 cells with gradient concentrations of BCL treatment for 48 h. Cells with blank medium treatment were set as a control. (c) Relative recycling of ANCA in NCM460 cells after treatment with the blank medium or 40 μM BCL for 48 h. The relative recycling was calculated with the amount recycled for ANCA compared to that of the control. (d) mRNA contents of p50, p65, IκB, and STAT1 in NCM460 cells after treatment with blank medium or 40 μM BCL for 48 h. (e) Protein contents of p50, p65, IκB, pIκB, STAT1, and pSTAT1 in NCM460 cells after treatment with blank medium or 40 μM BCL for 48 h. All data are expressed as the mean ± S.D. (n = 3 replicates/treatment). The protein expression of p50, p65, IκB, pIκB, STAT1, pSTAT1, and the protein/mRNA expression levels of FcRn were normalized to those of GAPDH or β-actin. *n.s* represents the differences that were not statistically significant (p > 0.05). */# represent p < 0.05; ** represents p < 0.01; *** represents p < 0.001 against the control.

### 3.3. BCL regulates FcRn via NF-κB signaling in NCM460 cells

After treatment with 40 μM BCL, we determined the expression of the p50/p65 heterodimer, IκB/pIκB, and STAT1/pSTAT1 to assess whether p50/p65 or STAT1-mediated signaling was involved in the regulation of FcRn by BCL. We found that BCL treatment downregulates both the mRNA and protein expression of p50, p65, and IκB in NCM460 cells; however, the expression of pIκB, STAT1, and pSTAT1 remains unchanged (Fig. 3 d-e). These results indicate that BCL may downregulate FcRn expression via the p50/p65 heterodimer.

To validate this discovery, NCM460 cells were first pretreated with siRNA targeting p50, p65, and STAT1 to suppress their expression, followed by treatment with 40 μM BCL. Upon treatment with the respective siRNA molecules, protein and mRNA levels of p50, p65, and STAT1 were notably reduced by approximately 80% and 95%, respectively, compared to the control (Fig. 4 b, c, e and g, h, j). Additionally, it was observed that FcRn protein and mRNA expression decreased by around 70% and 95%, respectively, in cells treated with p50/p65-targeted siRNA (Fig. 4 a and f), while there was an increase of nearly 80% and 200%, respectively, in cells treated with STAT1-targeted siRNA (Fig. 4 d and i).

**Fig. 4.**
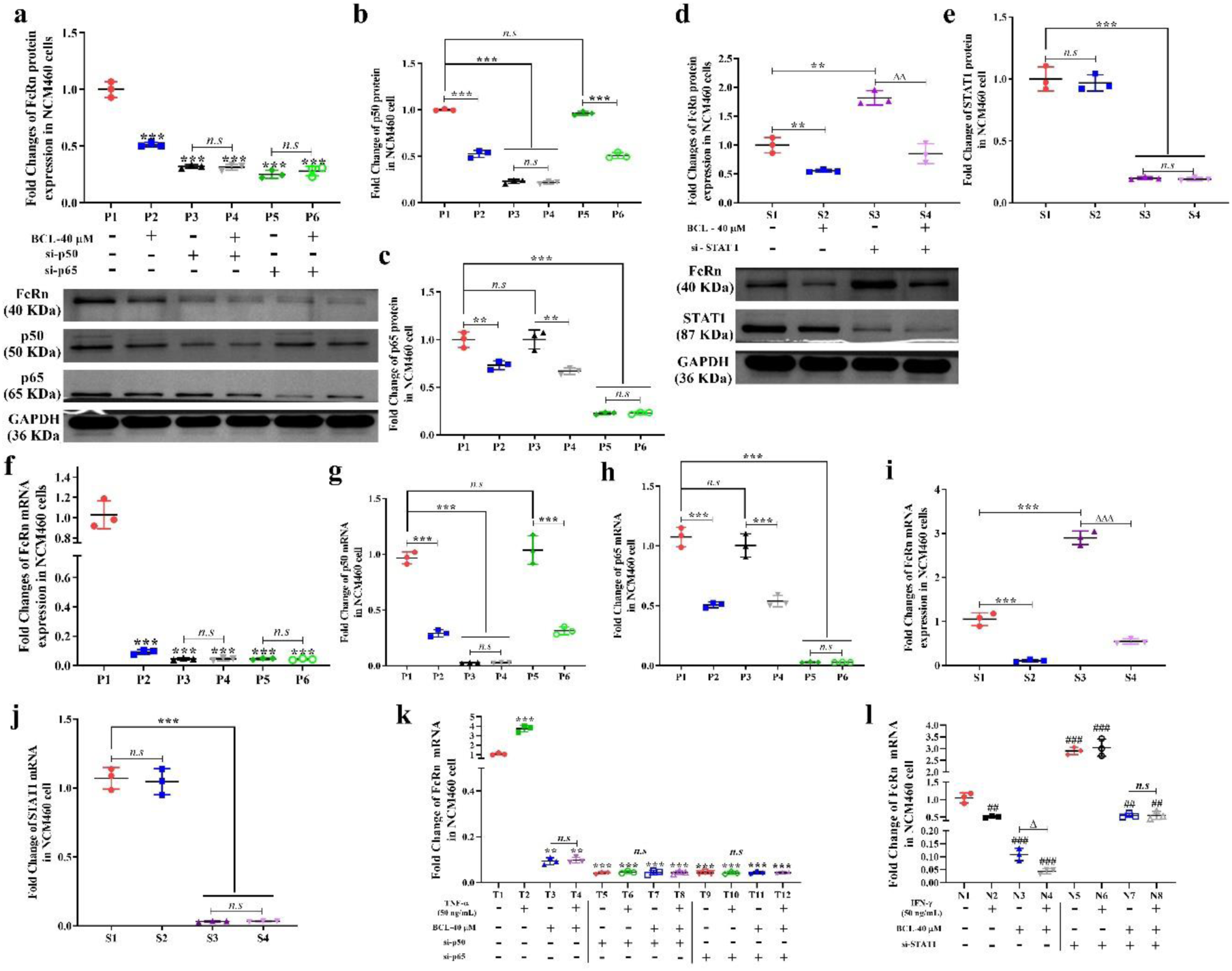
Effects of BCL on FcRn mRNA or protein expression in NCM460 cells with related transcription factors deficiency. (a–c) FcRn, p50, and p65 protein contents in NCM460 cells treated with blank medium or 40 μM BCL for 48 h in the presence or absence of p50 and p65-targeted siRNA. (d, e) FcRn and STAT1 protein contents in NCM460 cells treated with blank medium or 40 μM BCL for 48 h in the presence or absence of STAT1-targeted siRNA. (f–h) FcRn, p50, and p65 mRNA contents in NCM460 cells treated with blank medium or 40 μM BCL for 48 h in the presence or absence of p50 and p65-targeted siRNA. (i, j) FcRn and STAT1 mRNA contents in NCM460 cells treated with blank medium or 40 μM BCL for 48 h in the presence or absence of STAT1-targeted siRNA. (k) FcRn mRNA contents in NCM460 cells treated with blank medium or 40 μM BCL for 48 h in the presence or absence of p50 and p65-targeted siRNA with or without TNF-α (50 ng/mL) stimulation. (l) FcRn mRNA contents in NCM460 cells treated with blank medium or 40 μM BCL for 48 h in the presence or absence of STAT1-targeted siRNA with or without IFN-γ (50 ng/mL) stimulation. All data are expressed as the mean ± S.D. (n = 3 replicates/treatment). The mRNA or protein expression of FcRn, p50, p65, and STAT1 were normalized to those of GAPDH. Δ represents p < 0.05; **/##/ΔΔ represents p < 0.01; ***/###/ΔΔΔ represents p < 0.001; *n.s* represents the differences that were not statistically significant (p > 0.05).

Moreover, treatment with 40 μM BCL led to a reduction in FcRn protein and mRNA expression by about 50% and 90%, respectively, in cells pretreated with control or STAT1-targeted siRNA (Fig. 4 d and i). Conversely, cells pretreated with p50 and p65-targeted siRNA did not exhibit any inhibitory effects on FcRn mRNA and protein expression following additional BCL treatment (Fig. 4 a and f). Overall, these results suggest that BCL downregulating FcRn expression by diminishing the contents of the p50/p65 heterodimer in NCM460 cells.

### 3.4. BCL suppressed TNF-α activation of NF-κB signaling in NCM460 cells

As reported, TNF-α stimulation activated NF-κB signaling could up-regulate FcRn transcription level, while IFN-γ stimulation down-regulated FcRn transcription level via activating the JAK/STAT1 pathway[31, 32]. To further verify the role of these two signaling pathways played during the process of FcRn regulation by BCL, we treated NCM460 cells with blank medium or 40 μM BCL for 48 h in the presence or absence of p50, p65, or STAT1-targeted siRNA under conditions that with or without TNF-α or INF-γ stimulation. We found that TNF-α stimulation significantly increased the FcRn mRNA contents in NCM460 cells. This increase was dismissed when p50 or p65 deficiency, as well as BCL treatment (Fig. 4k). On the other side, IFN-γ stimulation reduced FcRn mRNA contents in NCM460 cells, and co-treated with BCL could further decrease it to a lower level (Fig. 4l). STAT1 deficiency eliminated the effects of IFN-γ stimulation on FcRn mRNA expression but didn’t impact FcRn mRNA reduction caused by BCL treatment (Fig. 4l). Collectively, we suggested that BCL may down-regulate FcRn expression via inhibiting of the p50/p65 heterodimer-mediated NF-κB signaling and STAT1-mediated signaling may not participate in the process of FcRn regulation by BCL.

### 3.5. BCL inhibits FcRn expression in UC mice via NF-κB signaling

Colonic FcRn mRNA and protein expression in the UC mice treated with BCL for 7 days was not altered. After 28 days of BCL administration, colonic FcRn mRNA and protein expression significantly decreased to approximately 80% and 50%, respectively, compared to the UC control. In contrast, SASP administration did not impact colonic FcRn expression during the 28-day treatment (Fig. 5a and 5b). Moreover, the colonic protein contents of p50 and p65 were decreased by about 60% and 80%, respectively, in UC mice after 28 days of BCL treatment (Figure 5c), but the expression of iκb/piκb and stat1/pstat1 remained almost unchanged compared to the UC control. These results were in line with those of the NCM460 cells, indicating that BCL downregulates colonic FcRn expression by inhibiting the p50/p65 heterodimer contents, which leads to the transcriptional reduction of FCGRT (the gene that encodes FcRn).

**Fig. 5.**
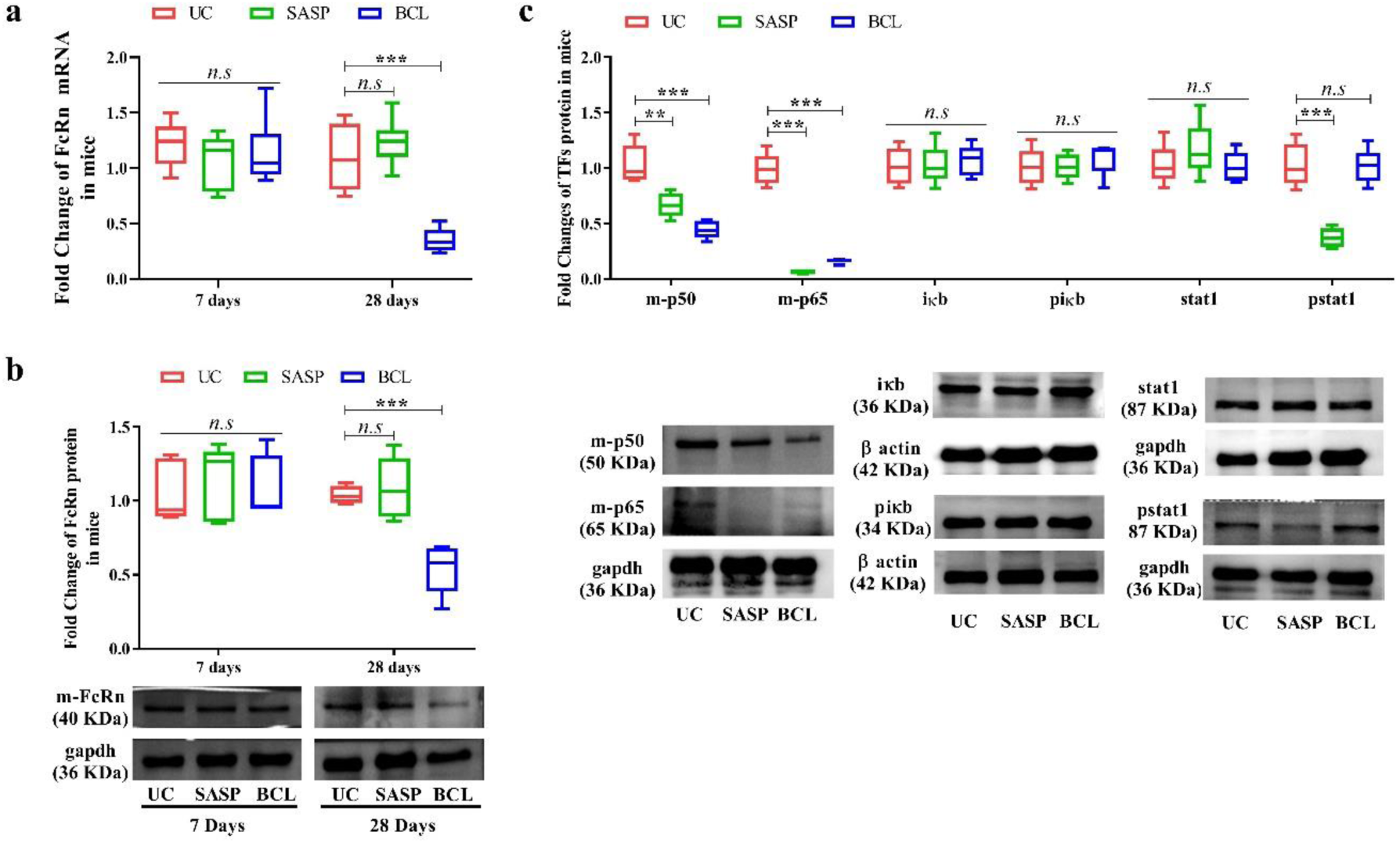
Effects of BCL on colonic FcRn and related transcription factor expression in mice with UC. (a, b) Colonic FcRn mRNA and protein contents in UC mice after SASP and BCL administration for 7 and 28 days. (c) Protein contents of colonic m-p50, m-p65, iκb, piκb, stat1, and pstat1 in UC mice after SASP and BCL administration for 28 days. All data are expressed as the mean ± S.D. (n = 6 replicates/treatment). Mice administrated with ultra-pure water were used as control (UC group). The protein expression of m-p50, m-p65, iκb, piκb, stat1, pstat1, and the protein/mRNA expression levels of FcRn were normalized to those of gapdh or β actin. * represents p < 0.05; ** represents p < 0.01; *** represents p < 0.001; *n.s* represents the differences that were not statistically significant (p > 0.05).

After 28 days of SASP administration, the p50, p65, and pstat1 protein contents were remarkably decreased, but the expression of iκb/piκb and stat1 were not altered. These results indicate that SASP administration simultaneously inhibits p50/p65 heterodimer and STAT1-mediated signaling. Since FcRn expression is increased by p50/p65 heterodimer stimulation and decreased by STAT1 signaling activation, we speculate that the SASP treatment suppresses both pathways, thus leading to the unaltered colonic FcRn expression after SASP treatment.

### 3.6. BCL reduced FcRn-mediated recycling and suppressed the ANCA-NET cycle in UC mice

BCL administration for 7 days did not alter the mouse serum IgG levels compared to the UC control. After 28 days of administration, the BCL-treated group significantly reduced serum IgG levels. SASP administration did not affect serum IgG levels at days 7, but slightly increased it at day 28. (Fig. 6a). These results are consistent with the changes in colonic FcRn expression listed above. Thus, we suggest that BCL-inhibited colonic FcRn expression impacts FcRn-mediated recycling in UC mice.

**Fig. 6.**
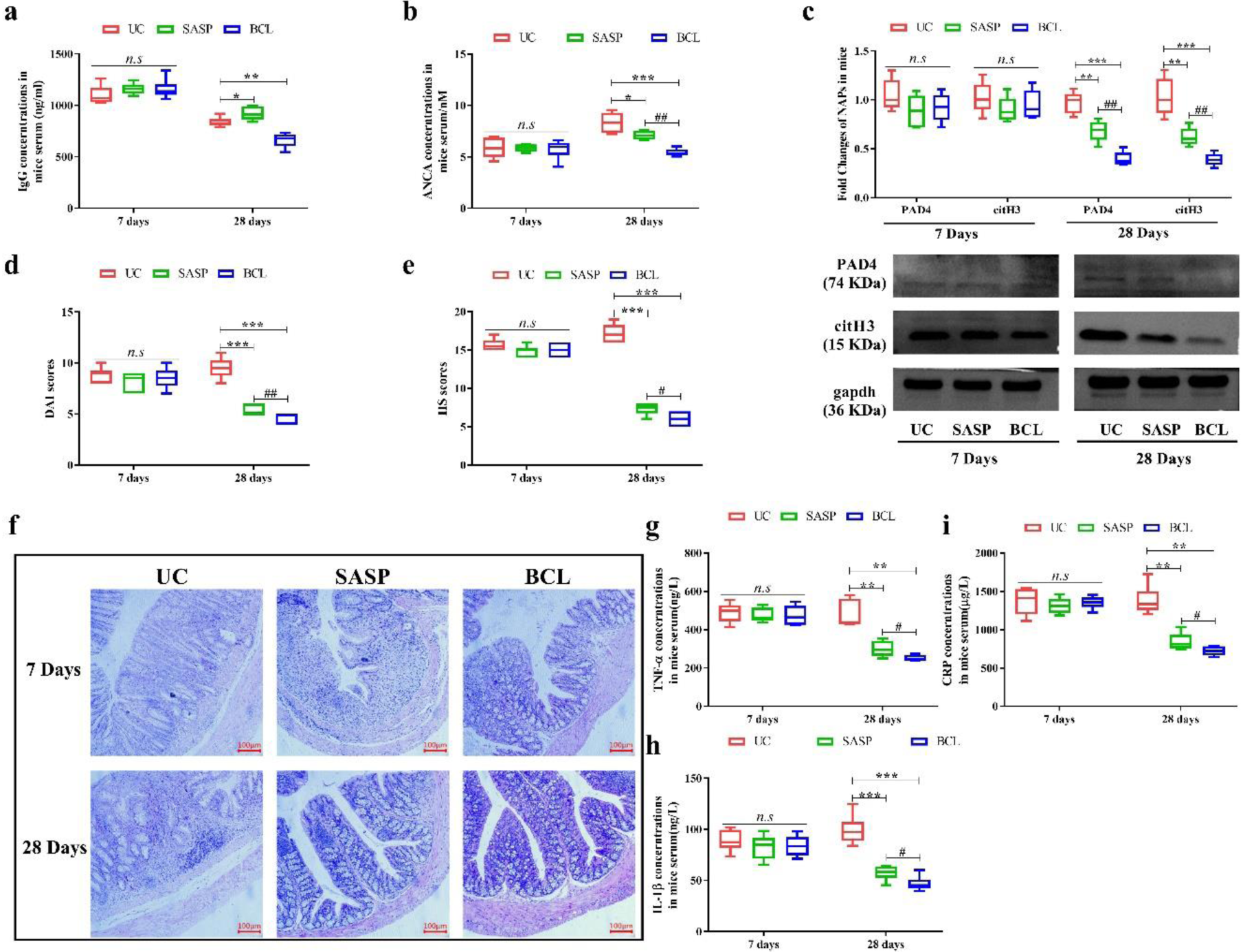
Effects of BCL on ANCA- NET cycle and UC relapse in mice. (a, b) Serum IgG and ANCA levels in UC mice after SASP and BCL administration for 7, and 28 days. (c) Colonic NAP (includes PAD4 and citH3) expression in UC mice after SASP and BCL administration for 7, and 28 days. The protein expression levels of PAD4 and citH3 were normalized to those of gapdh. (d, e) Calculation of DAI and HS in mice with UC after SASP and BCL administration for 7, and 28 days. (f) The representative histopathological images of mice colon in the control group (UC group), SASP group, and BCL group after 7, or 28 days of administration. (g–i) Serum concentrations of TNF-α, IL-1β, and CRP in UC mice after SASP and BCL administration for 7, and 28 days. All data are expressed as the mean ± S.D. (n = 6 replicates/treatment). Mice administrated with ultra-pure water were used as control (UC group). */# represents p < 0.05; **/## represents p < 0.01; and ***/### represents p < 0.001. *n.s* represents the differences that were not statistically significant (p > 0.05).

Both the BCL and SASP treatments had no impact on serum ANCA concentrations on day 7; however, they reduced the ANCA levels after 28 days, and BCL exhibited a greater reduction than SASP (Fig. 6b). Serum ANCA levels are associated with the inflammatory severity of UC; a reduction in serum ANCA levels implies a relief in inflammation, but FcRn inhibition may further increase ANCA clearance. This explains why the serum ANCA levels were reduced after 28 days of SASP treatment despite no change in colonic FcRn expression and FcRn-mediated recycling at that time, as well as why BCL exhibit stronger ANCA clearing effects than SASP as indicated by the day 28 findings. This finding is consistent with the colonic FcRn contents and FcRn-mediated recycling changes found earlier.

The colonic NAP contents remained unchanged on day 7, then decreased on days 28 in both the SASP and BCL groups compared with the UC group (Fig. 6c). Similar to the serum ANCA level results, BCL reduced it more than SASP by day 28. Taken together, our results suggest that a relatively short-term BCL administration of 7 days has no effect on serum ANCA levels and colon NET formation; but that long-term BCL treatment can inhibit colonic FcRn expression, reduce FcRn-mediated recycling, and accelerate ANCA clearance, which caused the greater suppression of the ANCA-NET cycle in the colons of UC mice found in this study.

### 3.7. BCL lowers UC recurrence based on FcRn blockade in UC mice

We further investigated whether BCL treatment lowered the recurrence of UC by determining DAI and HS scores since our results indicated that BCL inhibits FcRn expression and subsequently the ANCA-NET cycle. Interestingly, the DAI and HS scores (Fig. 6 d-e), serum IRI (TNF-α, IL-1β, and CRP) levels (Fig. 6 g-i), and colonic IRI (TNF-α and IL-1β) concentration (Fig. S4) remained unchanged after 7 days of treatment and were decreased on day 28 in both the SASP and BCL groups compared to the UC group. Similar to the serum ANCA results, BCL treatment decreased all indexes more than SASP did on day 28. These findings indicate that BCL is better able to reduce UC inflammation recurrency when administered long-term (i.e., 28 days) due to its inhibition of colonic FcRn expression and its greater suppression of the ANCA-NET cycle compared to SASP.

## 4. Discussion

Our previous study indicated that targeted FcRn inhibition remarkably ameliorates UC by suppressing the ANCA-NET cycle in rats with UC and is especially effective in withstanding UC relapse[38]. In the first part of this study, we once again confirm the efficacy of anti-FcRn therapy in mice with UC by downregulating of colonic FcRn expression with rAAV-Fcgrt-shRNA during the experimental-simulated inflammation progression and recurrence status. In the following works, we screened a series of compounds derived from Chinese traditional herbs to determine their effects on FcRn expression in HepG2 and HT-29 cells in vitro. We found that BCL significantly inhibits FcRn expression at nontoxic concentrations in both cell lines (data not shown). We verified this inhibition effect in the current study by first treating NCM460 cells with three nontoxic concentrations of BCL for 48 h, then administering BCL in UC mice for 28 days. In the former experiment, we found that BCL inhibits both the mRNA and protein contents of FcRn in cells with a dose-dependent manner. The results of the latter experiment further supported this finding since the BCL treatment markedly decreased the colonic FcRn mRNA and protein contents in the mice.

Considering that BCL’s inhibition effects on FcRn mRNA and protein expression were changed synchronously, we speculated that BCL might regulate FcRn expression by inhibiting FcRn’s transcription levels. Since previous studies have indicated that the transcription of FCGRT is primarily up- or downregulated by NF-κB elements (p50/p65 heterodimer) or STAT1[31, 32], we determined the expression levels of the p50/p65 heterodimer, IκB/pIκB, and STAT1/pSTAT1 to assess whether p50/p65 or STAT1-mediated signaling was involved in the regulation of FcRn by BCL. These results preliminarily indicate that BCL downregulates FcRn expression by reducing the contents of the p50/p65 heterodimer in NCM460 cells. When we pretreated NCM460 cells with specific siRNA molecules targeted against p50, p65, and STAT1, we found that BCL did not affect FcRn protein contents when the cells were deficient in p50 and p65. In contrast, BCL did downregulate the FcRn protein contents when the control siRNA molecule or STAT1-targeted siRNA molecules were used. Moreover, we used the known stimulus of NF-κB signaling, TNF-α, to activate this signaling pathway in NCM460 cells and found that BCL treatment could effectively dismiss TNF-α stimulation caused FcRn mRNA increase. Thus, we suggested that BCL did downregulate FcRn expression by inhibiting the p50/p65 heterodimer mediated NF-κB signaling. This inference is further supported by the data collected from the UC mice colons after 28 days of BCL administration.

Our data found that SASP administration for 28 days could inhibit both NF-κB and STAT1 signaling pathways in the colon of UC mice. The double inhibitions on these two pathways may have benefits on its anti-inflammatory effects. However, for FcRn transcriptional regulation, inhibition of NF-κB and STAT1 signaling showed the opposite results. Simultaneously inhibiting these two pathways may lead to a neutralized effect on FcRn expression. This speculation may partly explain that SASP administration didn’t affect colonic FcRn expression we observed. However, as the data showed, the inhibition extent of NF-κB signaling was much greater than STAT1 signaling by SASP, which means it may exit other mechanisms to neutralize FcRn down-regulation caused by NF-κB signaling inhibition in SASP-treated UC mice. R. B. Cejas *et al.* reported that DNA methylation in FCGRT could reduce its transcription[39]. Thus, we preliminarily detected the colonic DNA methyltransferase 1 (dnmt1) protein contents in UC mice after 28 day’s administration and found that SASP administration notably reduced colonic dnmt1 expression while BCL administration slightly increased dnmt1 expression (Supplementary data Fig. S5). The reduction of colonic dnmt1 contents by SASP could up-regulate FcRn transcription and may play a role in neutralizing FcRn down-regulation caused by NF-κB signaling inhibition. However, this speculation should be verified by further study. The NET formation has been proven vital in sustaining mucosal inflammation in UC[40]. ANCA, the specific biomarker of UC, can induce NET formation as well reported[10, 38, 41]. Moreover, ANCA can release lysozymes through capillaries, damage blood vessels and intestine tissues, and cause tissue damages through T cell-mediated cellular immune synergy in UC[10]. According to the published researches and our previous works, we suggested that ANCA may play an essential role in UC. Accelerating ANCA clearance from serum would benefit UC therapy and against UC relapse.

Above results indicated that BCL could inhibit FcRn expression via p50/p65 heterodimer mediated NF-κB signaling. In the following study, we found that BCL’s inhibition of FcRn further reduces ANCA recycling in vitro and accelerates the clearance of ANCA in UC mice, which results in an associated decrease in colonic NAPs and indicates diminished NET formation. Under inflammation relapse, long-term BCL treatment inhibits colonic FcRn and leads to a better therapeutic effect than SASP. This finding is likely due to a greater suppression of the ANCA-NET cycle in the mucosal lining of the colon under long-term BCL treatment. UC is a chronic, long-term condition that often includes periods of inflammatory progression, remission, and relapse with mild to severe symptoms during clinical practice. Anti-inflammation drugs, including traditional medications (e.g., SASP and 5-aminosalicylic acid) and targeted biological drugs (e.g., infliximab), remain the first choice in clinical UC treatment[43]. Although these drugs effectively reduce symptoms to a certain extent, they do not prevent inflammation relapse; therefore, inflammation relapse remains a great challenge in UC treatment[44]. BCL and its related formulations have been applied in UC treatment in China, but not as wide as the more established drugs. Given the long-term, recurring symptoms of UC and the results of this study, we suggest that BCL may serve as an additional beneficial treatment for UC patients in clinical practice. Our study reveals BCL’s novel therapeutic effects on UC recurrency from the perspective of immune regulation and contributes to the future application of BCL in UC treatment. However, the benefits of long-term BCL administration on UC recurrency and its potential side effects will need to be confirmed using further clinical data.

## 5. Conclusion

Our study confirmed the specific impact of downregulating FcRn in preventing UC relapse and indicates that BCL downregulates FcRn expression by inhibiting p50/p65 heterodimer-mediated NF-κB signaling, accelerating serum ANCA clearance, and reducing colonic NET formation. These effects exert therapeutic benefits against UC relapse. These findings may provide a preclinical foundation for a unique strategy in UC treatment to help lower the rate of UC recurrency and to improve general UC treatment.

## Abbreviations

ANCA: antineutrophil cytoplasm autoantibodies
anti-Fc-mAb: anti-rat Fc heavy chain heterodimer monoclonal antibodies
BCL: Baicalein
citH3: citrullinated histone H3
DSS: dextran sodium sulfate
DAI: disease activity index
FcRn: neonatal Fc receptor
HS: histopathological scores
IRI: inflammation-related index
IRP: inflammation recurrence period
NET: neutrophils extracellular traps
NAP: NET-associated protein
PAD4: protein arginine deiminase-4
rAAV: recombinant adeno-associated virus
SASP: Salazosulfapyridine
UC: Ulcerative colitis

## Declaration of Competing Interest

The authors declare that they have no known competing financial interests or personal relationships that could have appeared to influence the work reported in this paper.

## Funding

This work was supported by grants from the National Natural Science Foundation of China [grant number 82274319], and Postdoctoral Fellowship Program of China Postdoctoral Science Foundation [grant number GZC20231392].

## Acknowledgments

We would like to thank Enago (www.enago.cn) for English language editing.

## Author Contributions

Haoyang Hu: Investigation, Data Curation, Writing - Original Draft. Fuliang Lu: Investigation, Data Curation, Validation. Xudong Guan: Investigation, Formal analysis. Xuehua Jiang: Supervision, Writing - Review & Editing. Chengming Wen: Methodology, Investigation, Visualization, Writing - Original Draft. Ling Wang: Conceptualization, Supervision, Writing - Review & Editing.

## Data Availability Statement

The data underlying this article will be shared on reasonable request to the corresponding author.

---

**Fig. S1.**
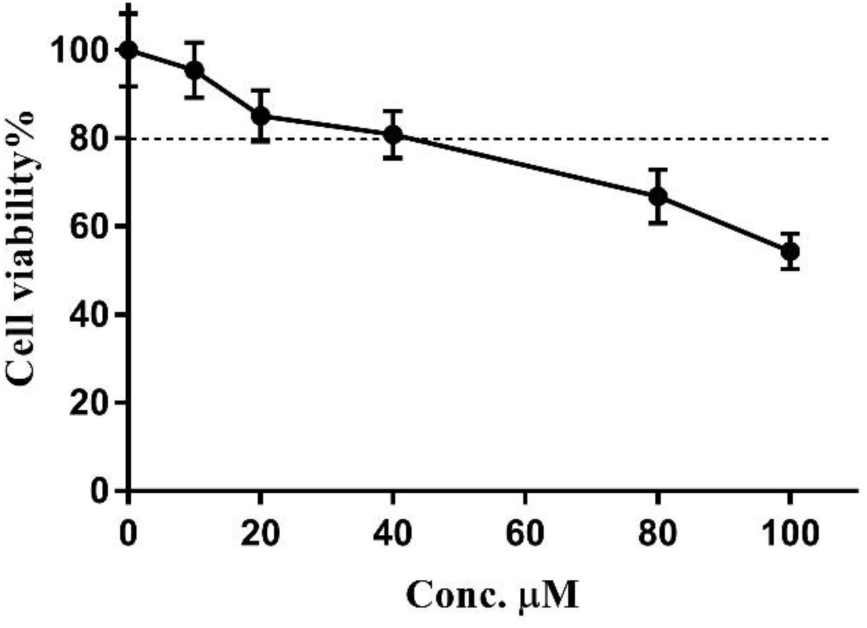
MTT assay of BCL on NCM460 cells. NCM460 cells were treated with the indicated concentrations of BCL (10, 20, 40, 80, or 100 μM) for 48 h, respectively. Data were expressed as the mean ± S.D. (n = 3 replicates/treatment).

**Fig. S2.**
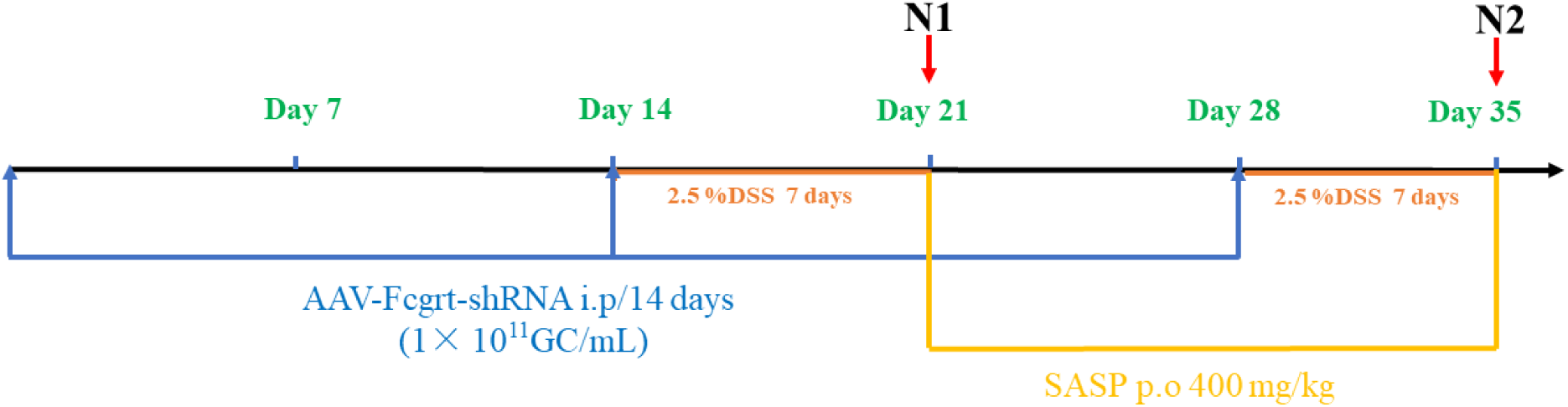
Process of animal experiments to verify the therapeutic effectiveness of anti-FcRn on UC. Male C57BL/6 mice weighing 25-30g were randomly divided into control group (12 mice), AAV injection group (12 mice), and SASP administration group (6 mice). In the AAV injection group, mice were intraperitoneally injected with rAAV-Fcgrt-shRNA (1×10^11^GC/mL) every 14 days. In the SASP administration group, mice were orally administered with SASP (400 mg/kg) starting from day 21. The control group was given ultra-pure (UP) water. Additionally, the control group and the SASP group mice were concomitantly intraperitoneally injected with shRNA-free rAAV vectors at the same frequency as the AAV group. All groups of mice were initially fed with UP water for 14 days, followed by a switch to 2.5% DSS solution for 7 days. They were then switched back to UP water on day 21 and further switched to 2.5% DSS solution for the last 7 days until the end of the experiment. On day 21, 6 mice from each of the control group and AAV injection group were euthanized (noted as N1, represented the inflammation progression period). On day 35, 6 mice from each of the control group, SASP administration group, and AAV injection group were euthanized (noted as N2, represented the inflammation recurrence period). Blood samples (500 μL) were collected from the retro-orbital plexus into microcentrifuge tubes. Then the mice were killed by cervical dislocation to separate the colons for biochemical and histopathological analyses.

**Fig. S3.**
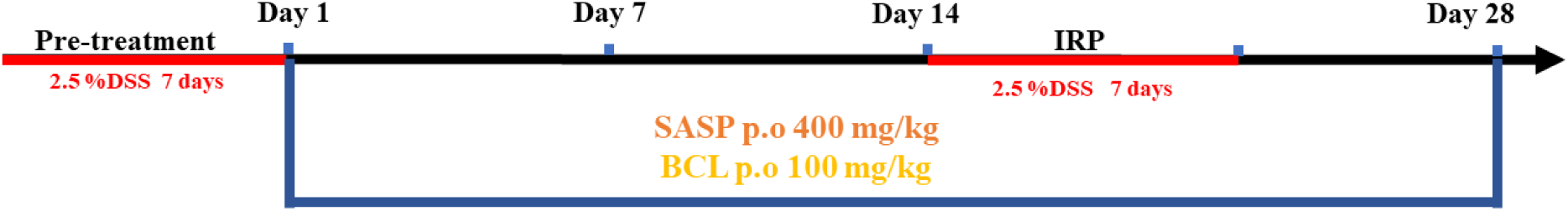
Process of animal experiments to evaluate the effectiveness of BCL treatment on UC relapse. Mice in group UC, SASP, and BCL were administered UP water, SASP (400 mg/kg, p.o/day), and BCL (100 mg/kg, p.o/day), respectively; Before the administration, mice in each group were first received 2.5 % (w/v) DSS for 7 days, followed by replacement with UP water for 14 days as a remission period. On the 22nd day, the 2.5 % (w/v) DSS solution was administered again for 7 days as an inflammation recurrence period (IRP) and then the 7-day remission again to the end of the experiment. During the experiment, six mice were sacrificed on Day 7 and Day 28. Blood samples (500 μL) were collected from the retro-orbital plexus into microcentrifuge tubes. Then the mice were killed by cervical dislocation to separate the colons for biochemical and histopathological analyses.

**Fig. S4.**
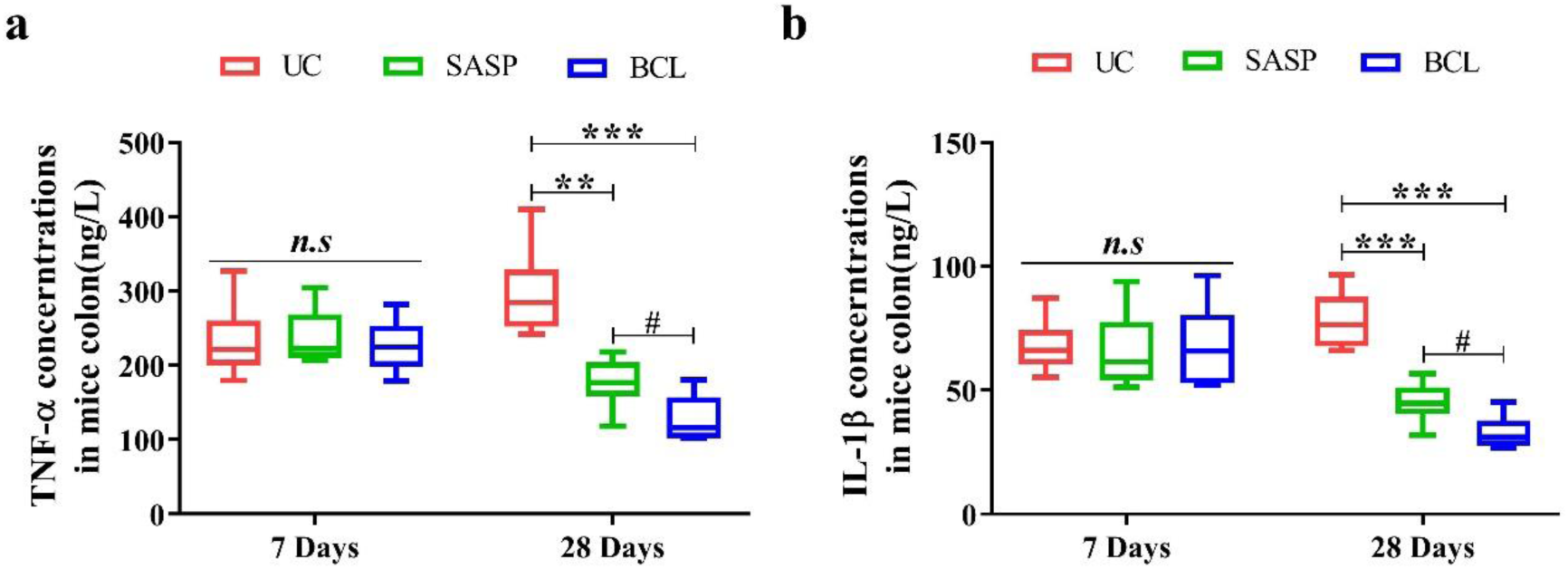
Colonic inflammation-related indexes determination in mice with UC. (a), (b) Colonic concentrations of TNF-α and IL-1β in UC mice after SASP and BCL administration for 7, and 28 days. All data are expressed as the mean ± S.D. (n = 6 replicates/treatment). Mice administrated with ultra-pure water were used as the control (UC group). */# represents p < 0.05; ##/** represents p < 0.01; and *** represents p < 0.001; *n.s* represents the differences that were not statistically significant (p > 0.05).

**Fig. S5.**
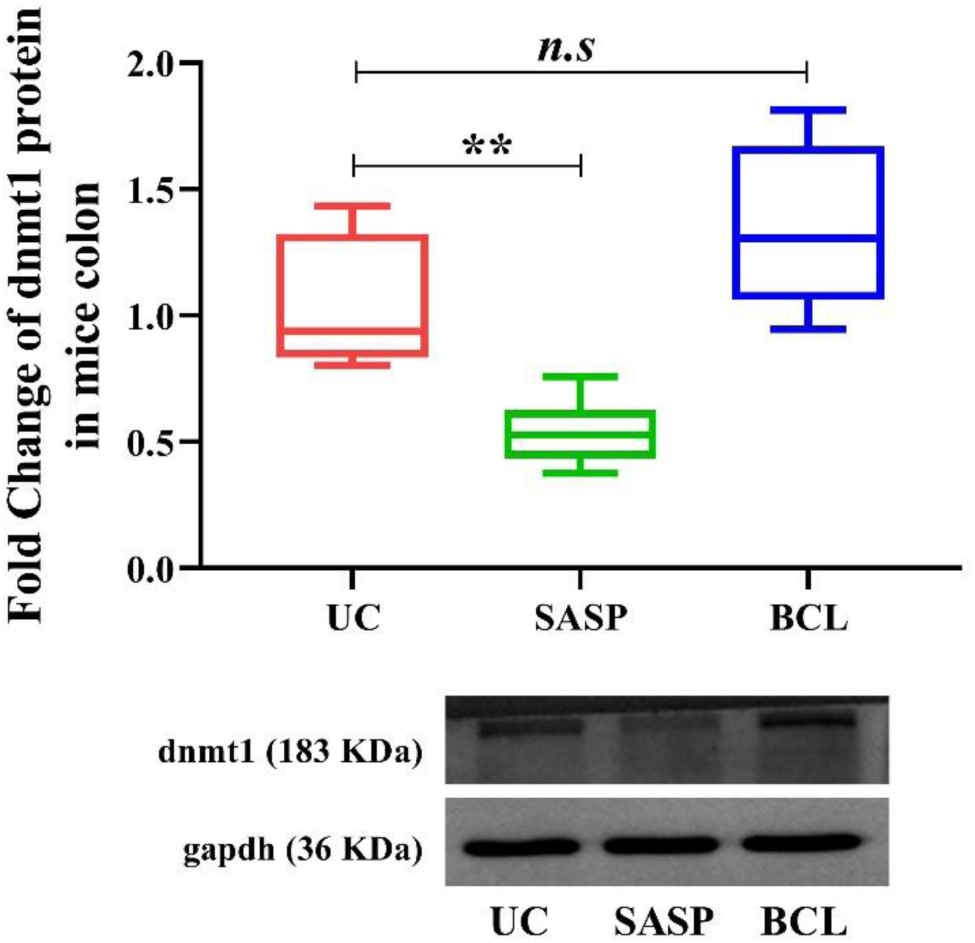
Colonic dnmt1 protein expression in UC mice after SASP and BCL administration for 28 days. Mice administrated with ultra-pure water were used as control (UC group). All data are expressed as the mean ± S.D. (n = 6 replicates/treatment). The protein expression of dnmt1 was normalized to those of gapdh. ** represents p < 0.01; n.s represents the differences that were not statistically significant (p > 0.05)

**Table S1.** Criteria for the colonic Disease activity index (DAI) scoring.

| Scores | 0 | 1 | 2 | 3 | 4 |
| --- | --- | --- | --- | --- | --- |
| body weight loss /% | 0 | 1~5 | 6~10 | 11~15 | >16 |
| stool consistency | Normal | Soft but shaped | Soft and unshaped | loose stool | — |
| occult blood /<br>Hematochezia | Normal | Occult blood<br>(+~+++) | Occult blood<br>(++++) | Hematochezia | Anal bleeding |

**Table S2.**
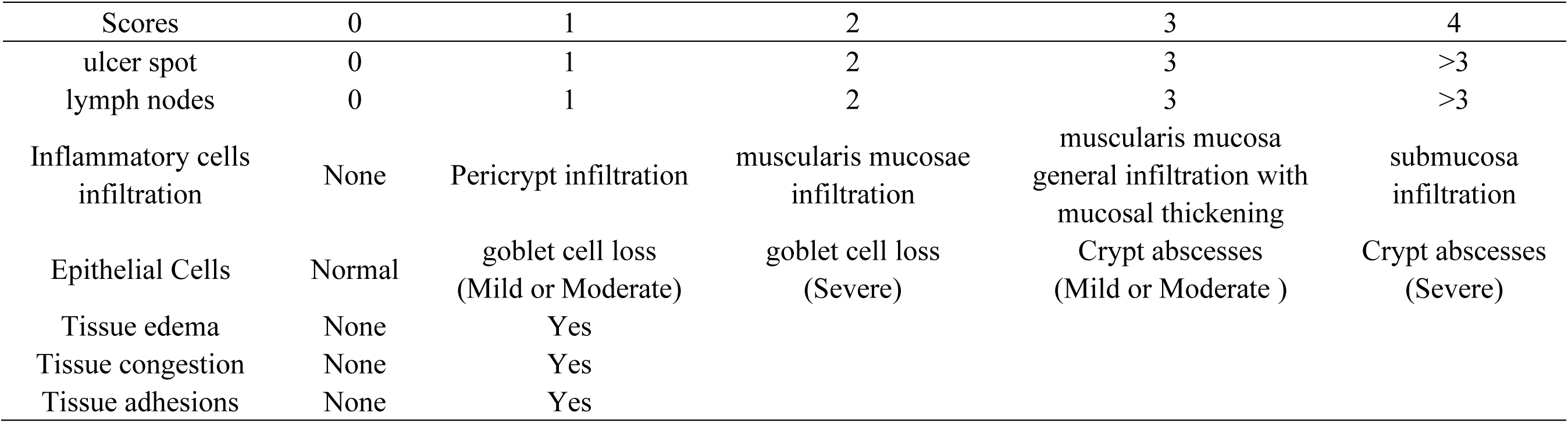
Criteria for the colonic Histopathological scoring (HS)

| Scores | 0 | 1 | 2 | 3 | 4 |
| --- | --- | --- | --- | --- | --- |
| ulcer spot | 0 | 1 | 2 | 3 | >3 |
| lymph nodes | 0 | 1 | 2 | 3 | >3 |
| Inflammatory cells<br>infiltration | None | Pericrypt infiltration | muscularis mucosae<br>infiltration | muscularis mucosa<br>general infiltration with<br>mucosal thickening | submucosa<br>infiltration |
| Epithelial Cells | Normal | goblet cell loss<br>(Mild or Moderate) | goblet cell loss<br>(Severe) | Crypt abscesses<br>(Mild or Moderate ) | Crypt abscesses<br>(Severe) |
| Tissue edema | None | Yes |  |  |  |
| Tissue congestion | None | Yes |  |  |  |
| Tissue adhesions | None | Yes |  |  |  |

**Table S3.**
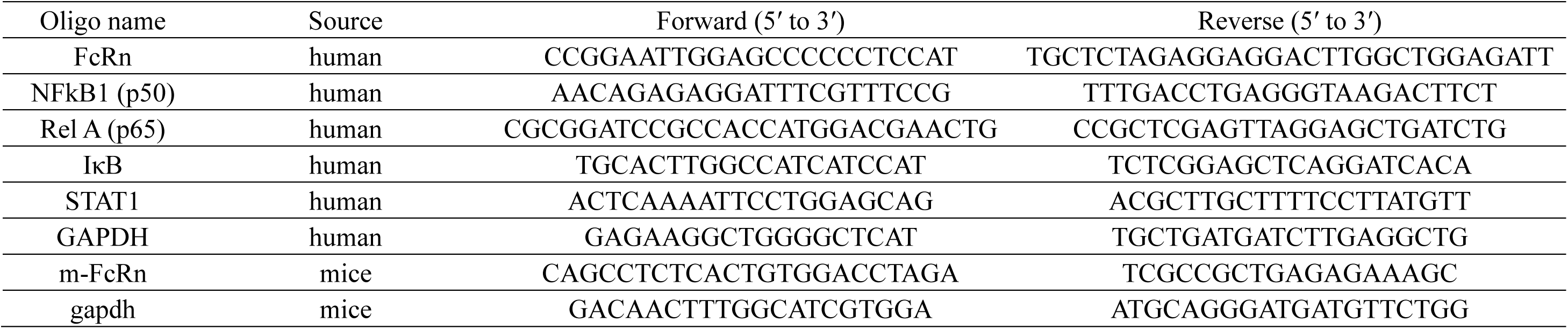
Oligonucleotides used in this study.

**Table S4.**
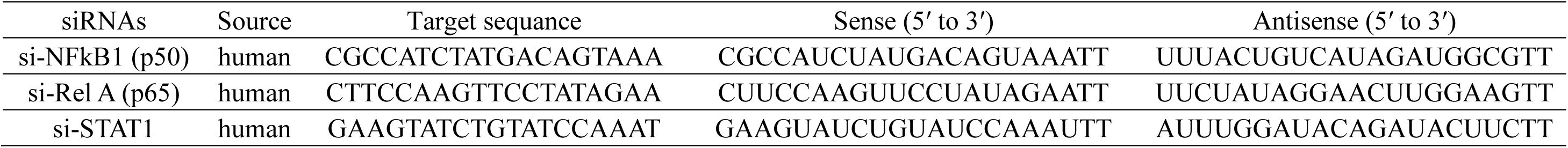
Sequences of siRNAs targeting p50, p65, and STAT1.

## Notes

### Competing Interest Statement

The authors have declared no competing interest.

